# Optimizing the Efficacy of Vaccine-Induced Immunotherapy in Melanomas

**DOI:** 10.1101/2025.01.06.631283

**Authors:** Ibrahim Chamseddine, Manoj Kambara, Priya Bhatt, Shari Pilon-Thomas, Katarzyna A. Rejniak

## Abstract

Cancer therapeutic vaccines are used to strengthen a patient’s own immune system by amplifying existing immune responses. Intralesional administration of the bacteria-based emm55 vaccine together with the PD1 checkpoint inhibitor produced a strong anti-tumor effect against the B16 melanoma murine model. However, it is not trivial to design an optimal order and frequency of injections for combination therapies. Here, we developed a coupled ordinary differential equations model calibrated to experimental data and used the mesh adaptive direct search method to optimize the treatment protocols of the emm55 vaccine and anti-PD1 combined therapy. This method determined that early consecutive vaccine injections combined with distributed anti-PD1 injections of decreasing separation time yielded the best tumor size reduction. The optimized protocols led to a twofold decrease in tumor area for the vaccine-alone treatment, and a fourfold decrease for the combined therapy. Our results reveal the tumor subpopulation dynamics in the optimal treatment condition, defining the path for efficacious treatment design. Similar computational frameworks can be applied to other tumors and other combination therapies to generate experimentally testable hypotheses in a fairly unrestricted and inexpensive setting.

## 1. Introduction

While the immune system provides the first line of defense against foreign bodies, such as viruses or cancer cells, the patient’s own activated T cells are rarely effective in killing large tumors. Thus, additional methods for boosting patients’ immune systems are desired. One of such methods is administering therapeutic cancer vaccines designed to eradicate cancer cells by strengthening a patient’s own immune system by either inducing new or amplifying existing immune responses [1-4]. When such vaccines are injected into a tumor, they transfect tumor cells and induce signals that activate the immune system response, resulting in increased tumor infiltration by the T cells.

Bacterial-based vaccines have been known for over a century to stimulate the immune system [5, 6]. One of the first studies of immunotherapy in the treatment of malignant tumors, which began in 1891 by William B. Coley, introduced streptococcal organisms into a cancer patient to stimulate the immune system [5]. Administration of attenuated mycobacteria, specifically Bacillus Calmette–Guérin (BCG) vaccine, is still successfully used in the clinical treatment of superficial bladder cancer [7, 8]. Studies have shown that anti-tumor immunity can be induced by activating TLRs in dendritic cells [9-11]. The addition of microbial extracts containing TLR ligands has been shown to induce tumor antigen presentation to T cells by activated dendritic cells, resulting in anti-tumor immunity [12, 13].

The cancer vaccine under consideration here is plasmid encoding the emm55 protein (emm55 vaccine), which is a serotyping protein normally expressed on the surface of the bacterium *Streptococcus pyogenes* [14, 15]. It has been shown that the lytic activity of *S. pyogenes* and a single application of live bacteria into established Panc02 tumors resulted in complete tumor regression [16]. This study also confirmed the leukocyte infiltration within the tumors with elevated IFN-γ production upon re-stimulation with tumor cells. This indicates that *S. pyogenes*, or its related proteins, are highly antigenic and may act as adjuvants for priming or activating tumor-specific T cells. In fact, the intralesional (IL) injection of plasmid DNA vaccine expressing emm55 induced a tumor-specific immune response and yielded clinical efficacy [14]. IL therapies have significant advantages as they can be used to treat tumors that are not resectable and allow avoidance of the off-target effects that are induced by standard therapies such as radiation and chemotherapy. Thus, IL treatment with therapeutic vaccines shows promise for strengthening a patient’s own immune systems to fight tumors.

However, the dynamics of interactions between native cytotoxic T cells, tumor cells, and therapeutic vaccine are quite complex. As a result of transfection of tumor cells by the vaccine, tumor-associated antigens are released that are captured by the dendritic cells (DCs), which, in turn, prime the T cells. The activated T cells infiltrate the tumor, bind to and kill the target cancer cells. This again leads to the release of specific antigens by the dying cancer, and this cancer-immune cycle is repeated [17]. However, the transfected tumor cells often have highly upregulated co-inhibitory ligands such as PD-L1, which may engage in binding to PD1 on T cells, resulting in exhaustion and deactivation.

The combination of emm55 cloned into a DNA plasmid vaccine and a monoclonal antagonistic antibody targeting PD1 was investigated in a murine model of melanoma [18]. A specific administration protocol consisting of one weekly injection of vaccine and two anti-PD1 injections per week was tested for three weeks, leading to significant tumor size reduction (∼70% smaller than the control case), but the tumor was not eradicated [18]. This was a motivation for our study to investigate more effective administration protocols using the mathematical optimization of and ordinary differential equations (ODE) model hierarchically calibrated with experimental data.

While general topics in the mathematical modeling of immunotherapy have been widely explored [19-23], the modeling of therapeutic cancer vaccines is still quite limited and includes the modeling of dendritic cell activation [24], the direct stimulation of T cells [25-28], and dendritic vaccines [29-31]. To our knowledge, the model presented here is the first to address the emm55 vaccine with the aim of finding the optimal treatment protocols. In general, various methods have been applied for scheduling anti-cancer treatments, including direct simulation studies [32-34], design optimization methods with fast convergence properties [35, 36], and optimal control methods for time-dependent objective functions [37, 38]. In our study, we use the Mesh Adaptive Direct Search (MADS) algorithm [39, 40], which has rigorous convergence properties and can handle a large number of optimization variables.

## 2. Methods

For this study, we have developed a continuous ordinary differential equations (ODE) model of native T cells interacting with tumor cells, exposed to a PD1 checkpoint blockade and a therapeutic emm55 vaccine. This in silico model was calibrated in a hierarchical fashion with experimental data on murine melanoma [18] and was used to predict the optimal administration protocols for this combination therapy. All of the computational methods used in our study are presented below.

### 2.1 Mathematical Model

The mathematical model describes temporal changes in the volumes of untransfected tumor cells (*U*), vaccine-transfected tumor cells (*I*), dead tumor cells (*W*), active infiltrating T cells (*T*), inactive/exhausted T cells (*J*), a therapeutic vaccine (*V*), and a PD1 checkpoint blockade (*A*). The vaccine is injected into the lesion at the schedule *u*_*v*_(*t*), cleared from the tumor at a rate *d*_*v*_, and induced transfection of tumor cells at a rate *k*_*v*_. The transfected tumor cells induced recruitment of the tumor-specific active T lymphocytes at a rate *k*_*T*_. Active T cells induced death of both types of tumor cells at a rate *c*_*W*._ However, T cell activity can be suppressed by the viable tumor cells at a rate *c*_*T*_, leading to T cell exhaustion. This process can be inhibited by anti-PD1 treatment that was injected at a schedule *u*_*A*_(*t*), cleared at a rate *d*_*A*_, and blocked PD1 receptors expressed on T cells at a rate *c*_*A*_. The viable tumor cells and active T cells can proliferate at rates *r* and *r*_*T*_, respectively. This model is presented graphically in the flowchart in Figure 1 and is formulated as a system of ordinary differential equations (1)-(7), where *N* = *U* + *I* + *W* + *T* + *J* is the total volume of tumor and immune cells.

**Figure 1:**
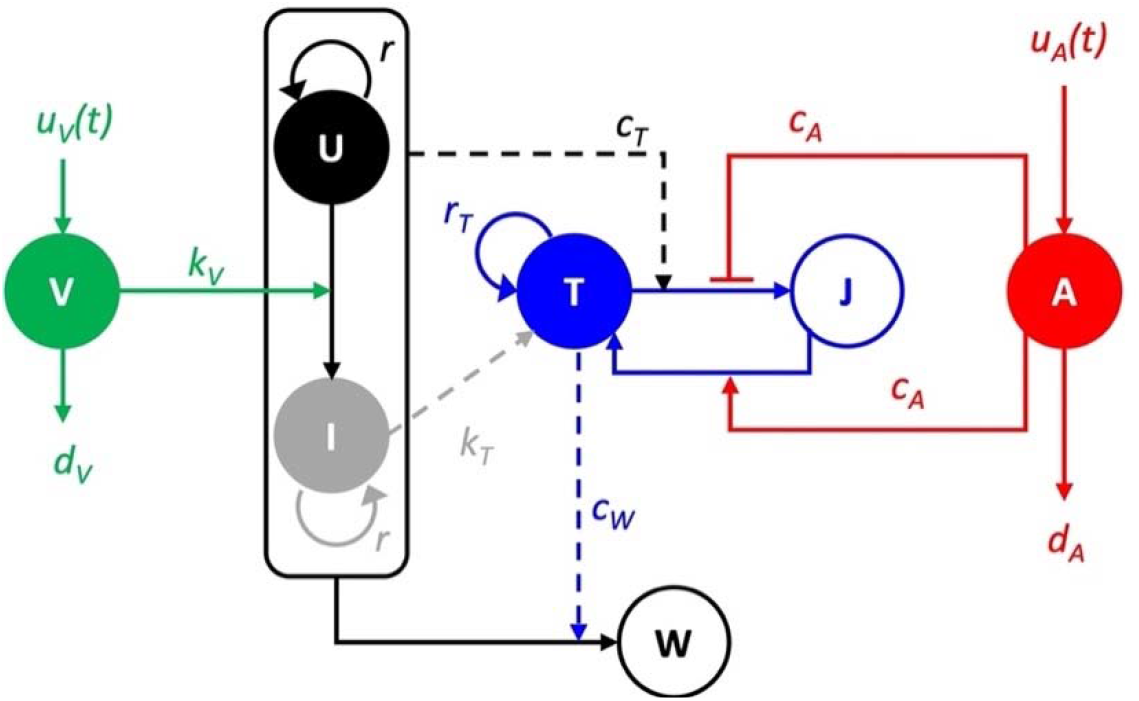
Mathematical model interaction diagram. Vaccine *V* is injected with a protocol *u*_*V*_(*t*) and transfects tumor cells *U*. The transfected tumor cells *I* recruit the specific T cells *T* that kill both untrasfected and transfected tumor cells generating dead cells W. T cells can become exhausted *J* as a result of blocking their PD1 receptors by tumor cells. Anti-PD1 checkpoint blockade *A* is injected with a protocol *u*_*A*_(*t*) and protects active T cells from being blocked, as well as reactivates the blocked T cells. In the diagram, filled circles refer to live or active cell populations, while hollow circles refer to dead or inactive cells; solid lines refer to the transfer between cell populations and dotted lines refer to feedback: either positive (arrows) or negative (flat ends).

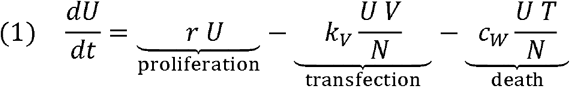

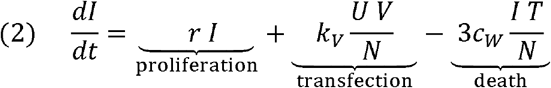

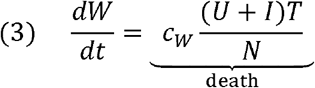

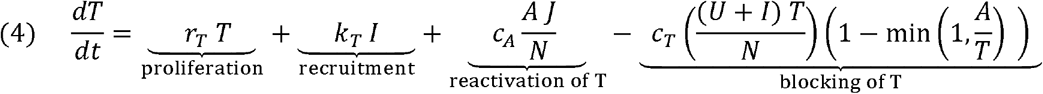

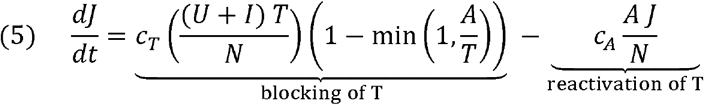

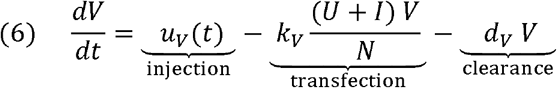

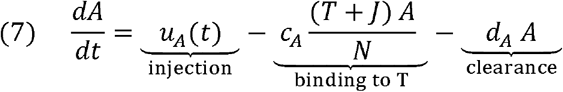

The initial tumor contains only untransfected tumor cells with the volume equal to the averaged experimental data—that is, *U*(0) = *U*_0_, *I*(0) = 0, *W*(0) = 0, *T*(0) = 0, *J*(0) = 0. Initially, there is no vaccine and no PD1 checkpoint inhibitor—that is, *V*(0) = 0 and *A*(0) = 0—then, their injections *u*_*V*_ and *u*_*A*_ followed the experimental schedule.

### 2.2 Optimization Procedures

Two types of optimization problems were solved here: calibration of model parameters to fit experimental data and scheduling of an efficient treatment combination.

For the calibration problem, the goal is to determine model parameter values *x* that minimize the least mean square error (*LMS*, equation (8)) between model outputs 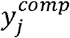 and experimental data 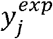 provided at *M* specific time points (*j* = 1… *M*).

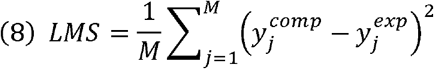

This may be subjected to specific constrains g_1_. The generalized form of the optimization problem formulated for model parameter calibration is shown in equation (9).

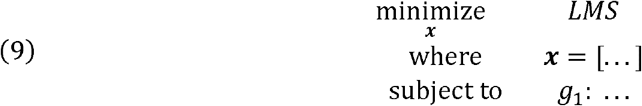

The entire model has nine parameters, as described in equations (1)-(7) that were calibrated in a hierarchical fashion. This means that the parameters fitted to a simpler model are fixed and carried in the more complex models. This process is described in detail in Section 3.1.

For the scheduling problem, the goal is to determine the injection schedule *x*_*V*_ that minimizes the tumor area at the end of treatment. The treatment doses are fixed and equal to the amount used in the experiments, so only the timing of the injections is optimized. The daily injections of vaccine or PD1 inhibitor are represented as binary vectors ***x***_*_ of cardinality equal to the treatment window (19 days). The vector element is equal to 1 if an injection is administered on the corresponding day and 0 if there is no injection on that day, with the exemption of weekend days, for which the vector elements are set to zero. This optimization problem may be subjected to specific constrains *g*_1_. The generalized form of optimization formulated for determining the optimal schedule is shown in equation (10), and the details are described in Sections 3.2 and 3.3.

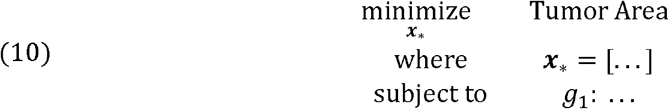

Both types of optimization problems were solved using the MADS approach. This derivative-free method does not require the evaluation of output gradients to proceed with the optimization search. This feature is useful for both types of optimization problems in this study. In the case of the calibration problem, it is expensive to compute the derivative of the *LMS* with respect to all model parameters at each iteration. In the case of the scheduling problem, the objective function (tumor area) is nondifferentiable with respect to the binary scheduling vectors. Therefore, the MADS algorithm was selected especially for these advantages and because of the rigorous convergence properties of this method. All optimization problems were solved using the NOMAD software package [41, 42] with MATLAB interface to provide communication between the optimization algorithm and the ODE model.

## 3. Results

To find optimal and experimentally feasible treatment schedules, we first calibrated the ODE mathematical model using previously published experimental data (Section 3.1). Next, the model was used to design optimal administration protocols for the vaccine alone (Section 3.2) and for a combination of the vaccine and the PD1 checkpoint blockade (Section 3.3). Finally, we compared the dynamics of all cell subpopulations for the optimized and experimental protocols (Section 3.4).

### 3.1 Mathematical Model Calibration to Baseline Experimental Data

The development of our mathematical model and current optimization studies were motivated by a recent publication showing the anti-tumor efficacy of the plasmid encoding emm55 vaccine in a murine model of B16 melanoma [18]. In that paper, mice were injected subcutaneously with B16 tumor cells and the tumor was allowed to grow for 7 days before the treatment started. Next, mice with palpable tumors received injections of the emm55 vaccine on days 7, 14, and 21. For combination therapy, mice also received injections of anti-PD1 blocking antibody twice per week, starting on day 8. Additionally, a control experiment with untreated tumors was performed over the same time period. Tumor measurements in all three experiments were recorded 2—3 times per week ([18], Figure 5D). The vaccine therapy yielded a 40% lower tumor area on day 25 compared to the control group, and the combined vaccine + anti-PD1 therapy resulted in a tumor area 69% smaller than in the control case. These longitudinal data, together with the final immunohistochemistry images ([18], Figure 2B), were used to hierarchically parameterize the ODE model equations (1)-(7).

**Figure 2:**
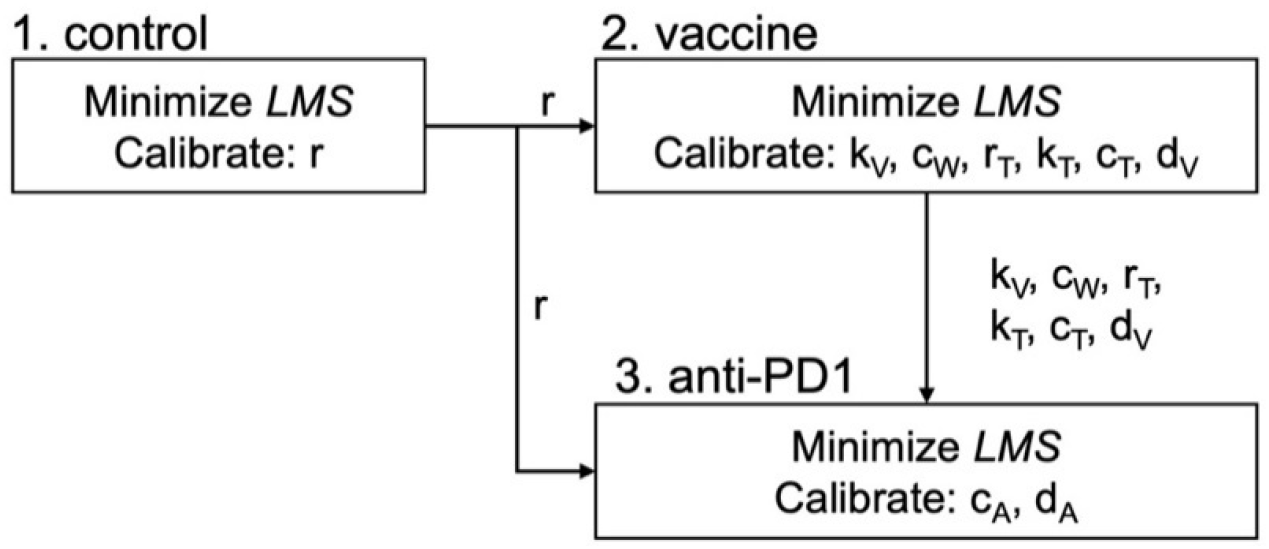
Hierarchical calibration framework. The entire model has been divided into three submodels: (1) the untreated tumor, (2) a tumor treated with the vaccine, and (3) a tumor treated with the combined vaccine and anti-PD1. These submodels are successively calibrated to corresponding experimental data of by minimizing *LMS*. The parameters calibrated in each submodel are listed in the corresponding boxes; the previously calibrated parameters listed along the arrows are kept constant and equal to the value calibrated in the previous submodel.

The hierarchical parameterization approach separates model parameters into groups that are appropriate for each set of experimental data. In this method, parameters calibrated using simpler models keep their fixed values in more complex models, and these models are used to calibrate the remaining parameters. This approach requires minimizing multiple *LMS* errors, however using a lower number of parameters at each calibration step in contrast to the case when all parameters are calibrated at once. A scheme of the hierarchical calibration framework is shown in Figure 2.

Due to the different units used in the experimental measurements (an injected emm55 vaccine is reported as volume and tumor sizes are reported as areas), we recalibrated all modeled quantities as volumes (mm^3^). However, to make comparisons with experimental data in optimization procedures, the total tumor volume was scaled back to the area of the tumor cross-section, assuming that the tumor had a spherical shape.

In the submodel 1 (Figure 2), we calibrated the tumor growth parameter *r* by fitting the equation (1) (with parameters *k*_*V*_ and *c*_*W*_ set to zero) to experimental data from an untreated mouse cohort. Experimental data were collected on days 7, 9, 11, 15, 18, 22, and 25 post the melanoma tumor initiation in the mice. We solved the following simple optimization problem:

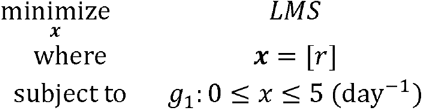

with the objective of finding a value of the parameter *r* that minimizes the *LMS* error between the exponential model of tumor growth and the average tumor size from mice in the control group, with number of data *M* = 7, and with the parameter *r* restricted to be between 0 and 5 (*g*_1_). This resulted in *r* = 0.28 day^-l^. This value of will be held fixed in all subsequent calibrations. The result of model calibration is shown in Figure 3A.

**Figure 3:**
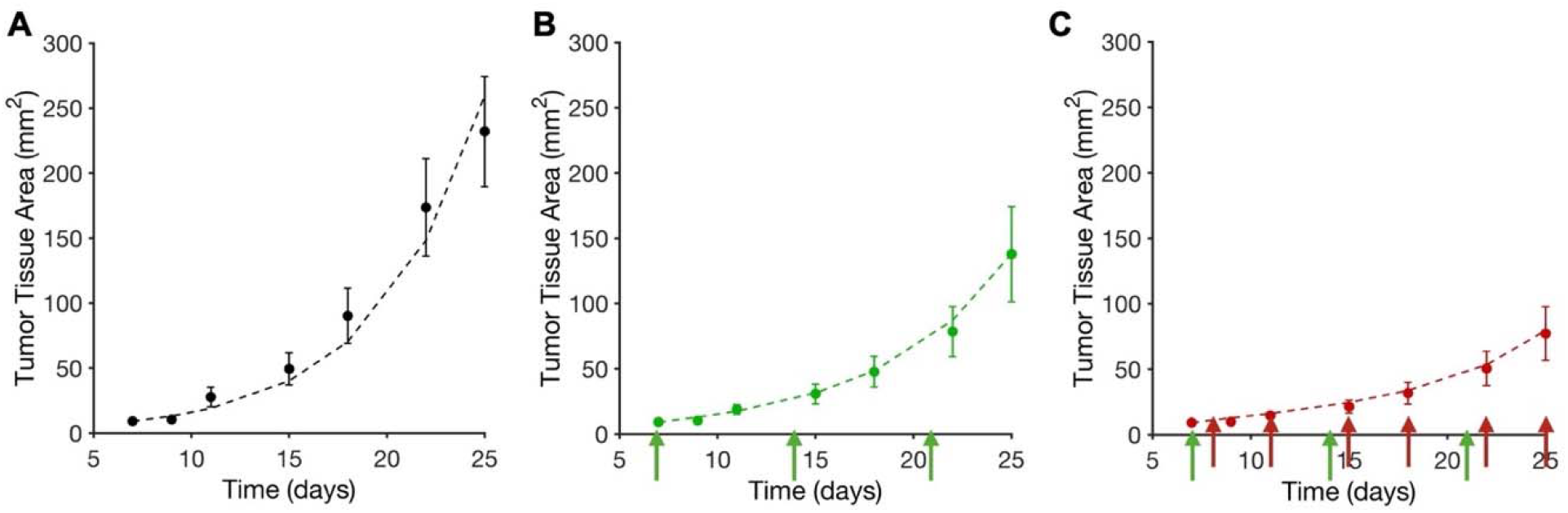
Model calibration with melanoma data. **A**. The tumor growth curve (dashed line) from submodel 1 was fitted to experimental data from an untreated mouse cohort (dots). **B**. The submodel 2 outcome (dashed line) was fitted to vaccine-treated mouse data (dots) with weekly injections of the emm55 vaccine indicated by arrows. **C**. The submodel 3 (full model) outcome (dashed line) was fitted to data from mice treated with combination therapy (dots); emm55 injection times are indicated by green arrows, and the anti-PD1 injections by read arrows. Error bars represent the standard error of the mean.

In submodel 2 (Figure 2), we used equations (1)-(6), with parameters *c*_*A*_ and *d*_*A*_ set to 0 and parameter *r* fixed to 0.28, as previously calibrated. The remaining six parameters were fitted to experimental data from a cohort of mice treated with the emm55 vaccine. This vaccine was injected weekly for three weeks, starting on day 7 post tumor installation. The parameter values were restricted to be between 0.01 and 5 (*g*_1_). According to the experimental results, the vaccine was able to transfect less than half of the tumor cells (*g*_2_) and at least 90% of the injected vaccine was cleared from the tumor microenvironment within 1 day of injection (*g*_3_). An additional constraint came from the analysis of histology images from untreated and vaccine-treated mice that were stained for T cell intensities ([18], Figure 2B). This yielded the percentage of tumor area occupied by T cells at the end of the treatment to be around 15.4% (*g*_4_) Thus, we solved the following optimization problem:

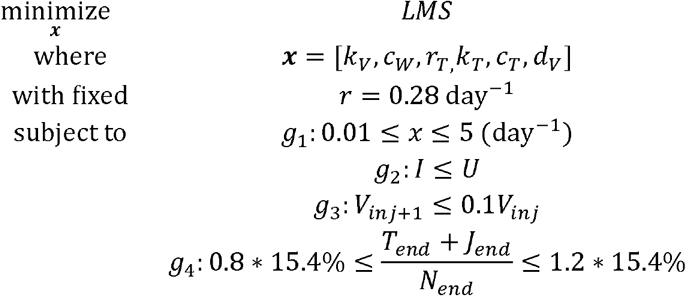

to find the values of the parameters *k*_*V*_, *c*_*w*_, *r*_*T*_, *k*_*T*_, *c*_*T*_, *d*_*V*_, which minimize the between the submodel 2 equations and the average tumor size from mice in the vaccine-treated group. The optimal values of all calibrated parameters are listed in Table 1. The results of the submodel calibration are shown in Figure 3B.

**Table 1:** Calibrated model parameters. Parameter values from equations (1)-(7) were calibrated in the hierarchical way (Figure 2), with those calibrated in the indicated submodel shown in bold, and other parameters fixed to the values calibrated in the previous model(s).

| Parameter and Description |  | submodel 1 | submodel 2 | submodel 3 |
| --- | --- | --- | --- | --- |
| $r$ | Proliferation rate of tumor cells (day<sup>-1</sup>) | **0.28** | 0.28 | 0.28 |
| $k_V$ | Transfection rate of tumor cells (day<sup>-1</sup>) | - | **0.49** | 0.49 |
| $c_W$ | Rate of tumor cell death by t cells (day<sup>-1</sup>) | - | **5.00** | 5.00 |
| $r_T$ | Proliferation rate of T cells (day<sup>-1</sup>) | - | **0.01** | 0.01 |
| $k_T$ | Rate of T cell recruitment (day<sup>-1</sup>) | - | **0.13** | 0.13 |
| $c_T$ | Rate of T cell blocking by PD1 (day<sup>-1</sup>) | - | **1.31** | 1.31 |
| $d_V$ | Clearance rate of vaccine (day<sup>-1</sup>) | - | **3.94** | 3.94 |
| $c_A$ | Rate of PD1 inhibition | - | - | **1.06** |
| $d_A$ | Clearance rate of anti-PD1 (day<sup>-1</sup>) | - | - | **1.83** |

In submodel 3 (Figure 2), we used the full set of equations (1)-(7) to calibrate the remaining parameters, *c*_*A*_ and *d*_*A*_, by fitting the model outcome to the experimental data from a cohort of mice treated with a combination of the emm55 vaccine and a PD1 checkpoint inhibitor. As before, the vaccine was injected weekly for three weeks and anti-PD1 twice weekly starting on day 8. All other model parameters were fixed to the previously calibrated values (compare Table 1). The only optimization constraint was to restrict the parameter values to be between 0.01 and 5 (*g*_1_). Thus, we solved the following optimization problem:

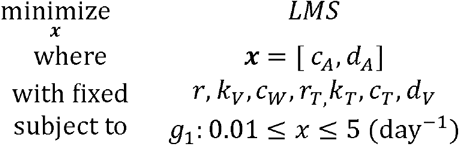

The optimal values of all calibrated parameters are listed in Table 1. The results of the model calibration are shown in Figure 3C.

### 3.2 Optimizing the Vaccine Administration Protocol

The emm55 vaccine treatment was used to enhance infiltration of cytotoxic lymphocytes into the tumor and thus to increase T cell-induced death of tumor cells. While the experimental vaccine injection protocol resulted in the diminished tumor size (Figure 3B) in comparison to the untreated tumor (Figure 3A), the extent of tumor reduction may depend on the timing and frequency of emm55 treatment. In this section, we used the MABS optimization method to determine more efficient vaccination protocols.

The experimental protocol for the vaccine treatment was a single weekly injection for 3 weeks (protocol V1 in Figure 4). This treatment reduced the total tumor area to 59% of that in the untreated group (Figure 3A). To search for a more efficient vaccination protocol, we used mathematical optimization with an objective function to minimize the tumor area at the end of the treatment period (i.e., at day 25). The optimization problem was formulized as follows:

**Figure 4:**
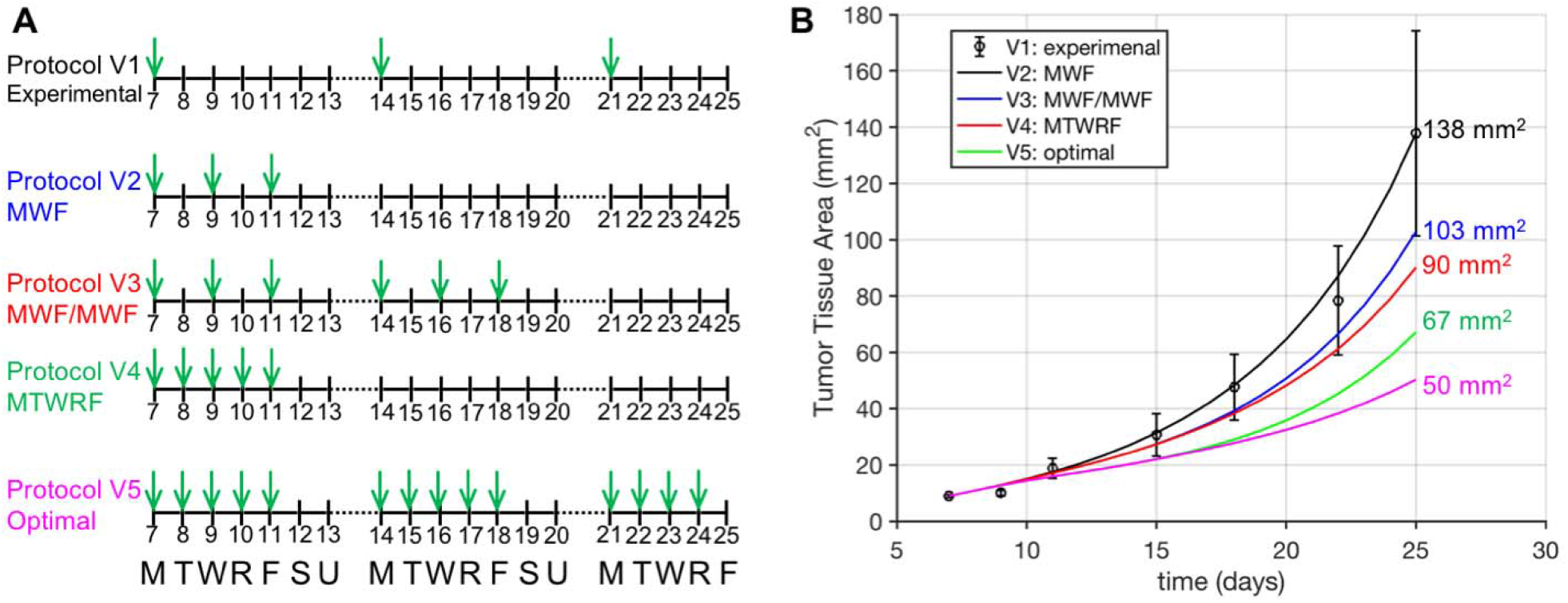
Comparison of vaccine protocols designed by computational optimization. **A**. Timelines of five protocols with arrows representing vaccine injections. The numbers represent the days after melanoma initiation in the mouse model. All treatments start on day 7. Injections are only performed on weekdays (M, T, W, R, and F), and no injections are done on weekends (S and U). **B**. Tumor growth curves corresponding to each of the five protocols (V1-V5). The final tumor areas on day 25 are listed in the color corresponding to the protocol. The black curve represents the experimental protocol, with error bars illustrating the standard mean of the error.

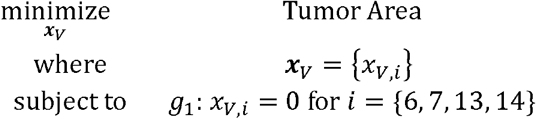

where *x*_*V*_ = {*x*_*V,i*_} is a binary variable with length 19 corresponding to the number of possible treatment days, for which values were set to 1 on the days the vaccine injections were scheduled or 0 if the injections were not administered. However, no injections took place on weekends, therefore, the elements of *x*_*V*_ that fell on weekend days were set to zero—this condition is included as constraint *g*_1_ in the above formulation. Thus, the design space for vaccine injections is composed of 2^15^ candidate protocols.

The employed MADS optimization method identified the optimal protocol (V5 in Figure 4), which consists of daily injections (except weekends and the last day of the protocol). This protocol resulted in a reduction of tumor area to 36% of that in the experimental protocol (V1 in Figure 4). However, such frequent vaccine administration may induce a toxic risk or undesirable immune responses; thus we investigated an additional set of protocols in an attempt to find near-optimal ones but with less frequent injections, and thus with less toxicity. These are presented in Figure 4A and summarized in Table 2.

**Table 2:**
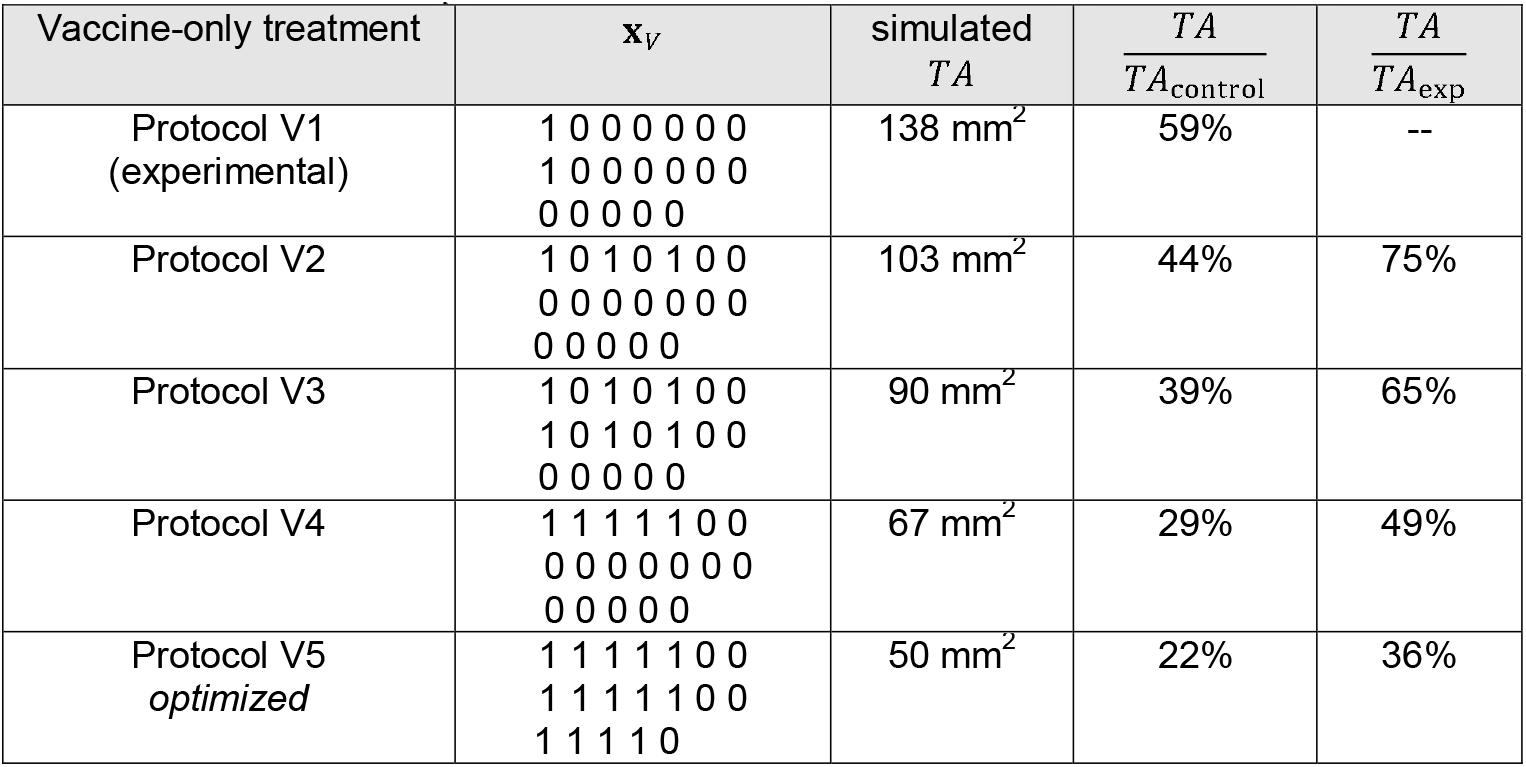
Results of the simulated vaccine protocols. For each of the five protocols (V1-V5), the corresponding 3-week schedule *x*_*V*_ is shown together with final simulated tumor area (*TA*), tumor area in relation to an untreated tumor (*TA/TA*_*control*_), and tumor area in relation to the experimental protocol (*TA/TA*_*exp*_) to show area reduction in the simulated schedules.

First, we examined the effect of a protocol with the same number of injections as the experimental protocol but with all injections scheduled within the first week (Monday, Wednesday, Friday; protocol V2 in Figure 4). This alteration of the baseline protocol reduced the tumor area to 75% of that from the experimental protocol (V1) and 44% of the size of the untreated tumor (Table 2). Next, we examined an extension of protocol V2 by continuing this schedule during the second week (protocol V3). Although the number of injections doubled, the reduction in tumor area only improved by 10% (compare protocols V2 and V3 in Table 2). This shows that most of the tumor reduction occurred in response to the vaccine injected during the first week. This conclusion is further emphasized by comparing protocols V3 and V4. Protocol V4 has one less injection than protocol V3, but all injections are administered during the first week (Figure 4), resulting in a significantly reduced tumor area at day 25 (49% vs. 65% of the tumor area in the untreated case; compare Table 2). This suggests that protocols with early repetitive vaccine injections are more effective and that adding more injections later has a limited effect. Moreover, the performance of protocol V4 was similar to the optimal protocol V5 when they were both compared to the final size of the untreated tumor (control case). There was only a 7% improvement in V5 vs. V4 (Table 2); however, schedule V5 required more injections over a longer time than protocol V4.

### 3.3 Optimizing Anti-PD1 Administration Protocols

The anti-PD1 treatment was used to block binding between tumor cells and T cells, and thus to enable more effective killing of tumor cells by T cells. The experimental protocol combining the emm55 vaccine and anti-PD1 treatment resulted in a significant reduction in tumor size (Figure 2C) when compared to the untreated tumor (Figure 2A). However, this final tumor was larger than the tumors predicted by our two best, optimized vaccination protocols. Therefore, in this section, we used optimization methods to determine a more efficient protocol that combines vaccination and anti-PD1.

The experimental protocol for the combined emm55 vaccine and anti-PD1 treatment was a single weekly vaccine injection (Mondays) accompanied by two weekly anti-PD1 injections (Tuesdays and Fridays) for 3 weeks (protocol VA1 in Figure 5). This protocol reduced the tumor area to 34% of that in the untreated group (Figure 3A) and to 50% of that in the vaccine-treated group without anti-PD1 treatment (Figure 3B). It is, however, possible that the different timings of both injections may have a therapeutic benefit. To search for a more efficient combination protocols, we considered the three vaccine protocols from Section 3.2 (V2−V4), and optimized the anti-PD1 protocol for each case, with an objective function to minimize tumor area at the end of the treatment period (i.e., at day 25). The optimization problem was formulized as follows:

**Figure 5:**
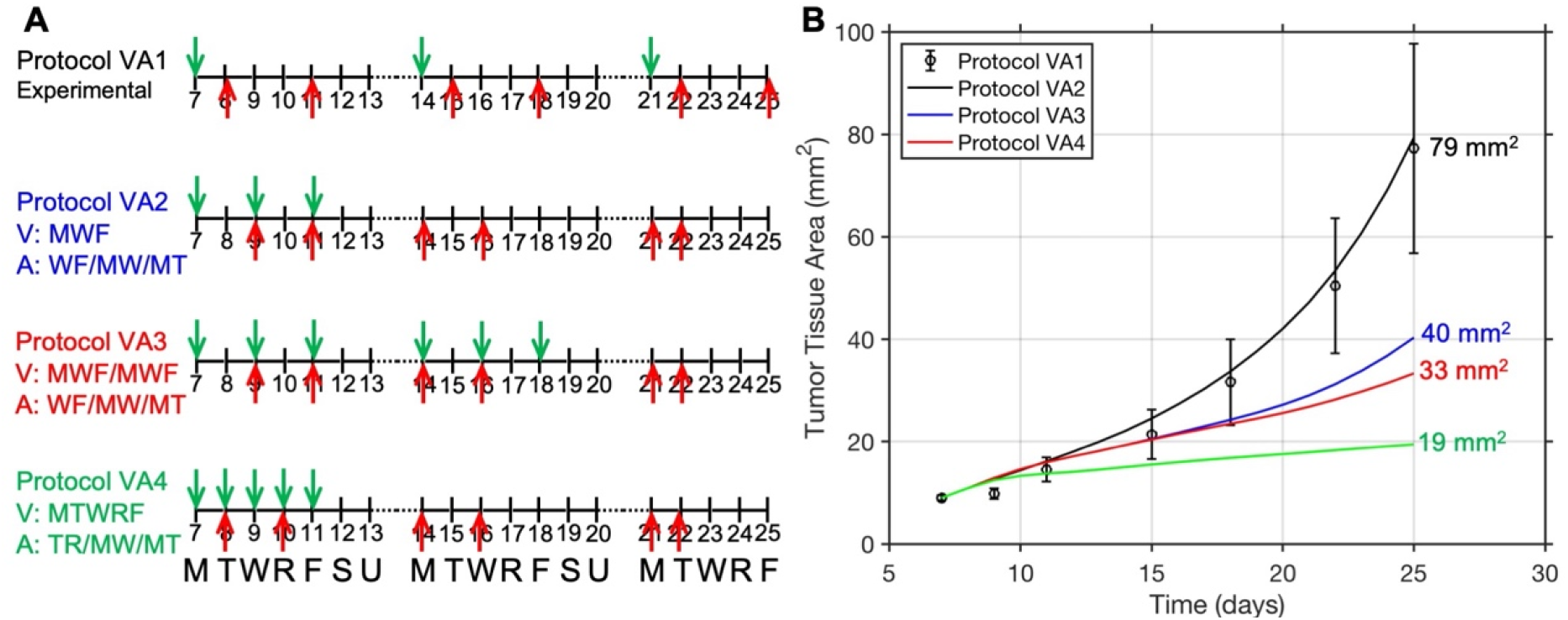
Optimized anti-PD1 protocols. **A**. Timelines of four protocols with arrows representing injections of a vaccine (green) and anti-PD1 (red). The vaccine protocols were previously designed (V1−V4) and held fixed here. The anti-PD1 protocols were optimized for each vaccine schedule. The numbers represent the days after melanoma initiation in the mouse model. All treatments started on day 7. Injections were administered only on weekdays (M, T, W, R, and F), and no injections were done on weekends (S and U). **B**. Tumor growth curves corresponding to each of the four protocols: VA1−VA4. The final tumor areas on day 25 are listed in the corresponding colors. The black curve represents the experimental protocols with error bars illustrating the standard mean of the error.

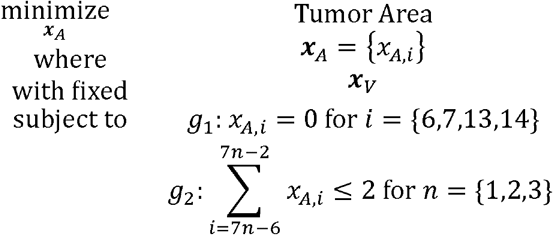

where, vector x_*A*_ = {x_*A,i*_} is a binary variable of length 19 representing the duration of treatment. An element x_*A,i*_ was set up to 0 if no anti-PD1 injection was scheduled on day *i*, and was equal to 1 otherwise. Again, no injections were scheduled on the weekends (constraint *g*_1_). In addition, a maximum of tqo anti-PD1 injections were allowed per week due to toxicity concerns (constraint *g*_2_). For each fixed vaccination schedule x_*V*_ (for protocols V2−V4), the MADS optimization method was used to optimize the corresponding schedule x_*A*_ (protocols VA2−VA4). This is presented in Figure 5 and summarized in Table 3.

**Table 3:**
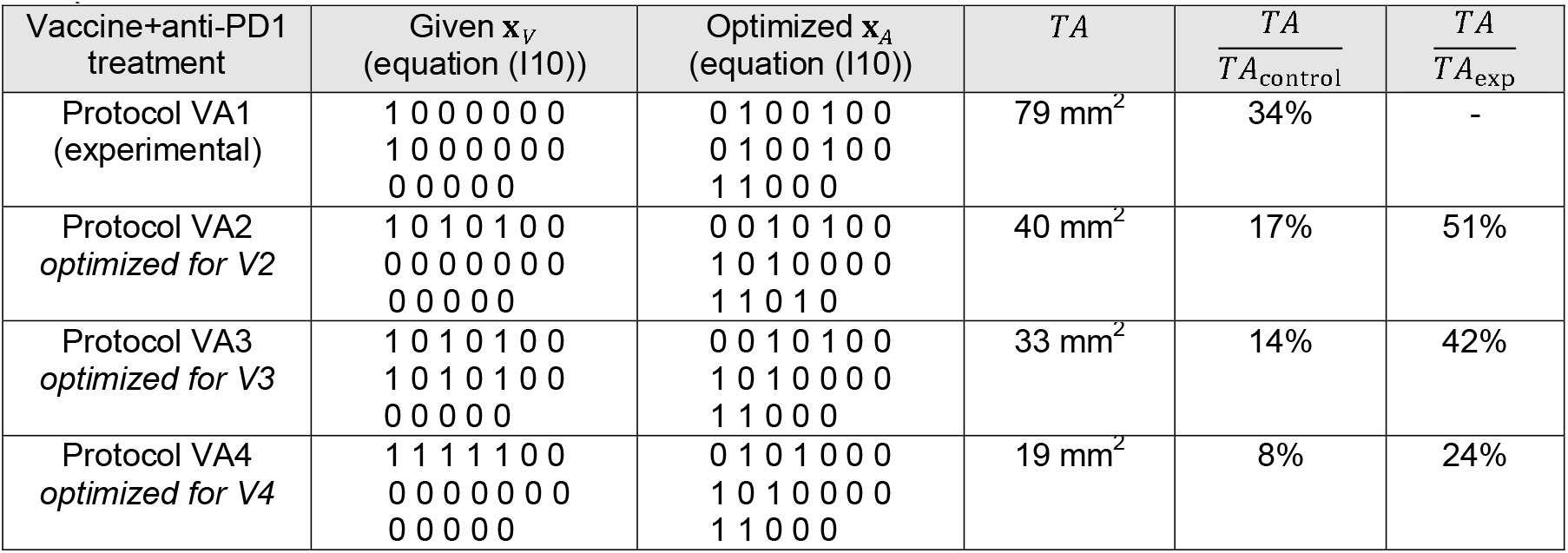
Results of simulated combinations of vaccine and anti-PD1 protocols. For each of the four protocols (VA1−VA4), the corresponding 3-week schedule of vaccine *x*_*V*_ and anti-PD1 *x*_*A*_ are shown together with the final simulated tumor area (*TA*), tumor area in relation to an untreated tumor (*TA/TA*_*control*_), and tumor area in relation to the experimental protocol (*TA/ TA*_*exp*_) to show area reduction in the simulated schedules.

The first optimized schedule of anti-PD1 treatment was performed for vaccine schedule V2, in which all vaccine injections were scheduled in the first week. The employed MADS optimization method identified the optimal protocol (VA2 in Figure 5), which consisted of irregularly scheduled anti-PD1 injections on Wednesday and Friday of the first week, then on Monday and Wednesday of the second week, and finally on Monday and Tuesday of the last week. Both protocols (VA2 and VA1) consist of the same number of injections (vaccine and checkpoint inhibitor) but differ in the details of the schedule. In protocol VA2, all vaccine injections are scheduled in the first week only, and the first anti-PD1 injection is delayed to allow for an initial T cell expansion. The subsequent two anti-PD1 injections are meant to boost T cells’ response. This protocol resulted in a half reduction of tumor area when compared to the experimental protocol for combination therapy (VA1 in Figure 5 and Table 3).

Next, we considered the case in which three more vaccine injections were added in the second week (protocol V3). The MADS optimization method determined the optimal anti-PD1 schedule (protocols VA3 in Figure 5), which, surprisingly, was not different from the previous case (protocol V2). However, the addition of these three vaccine injections in week 2 enhanced tumor reduction as compared to protocol VA2, but this improvement was minimal (Table 3). Finally, we considered vaccination protocol V4, which consists of five daily vaccine injections in the first week only. With this schedule fixed, we optimized the anti-PD1 injection protocol (VA4 in Figure 5), which resulted in first-week injections on Tuesday and Thursday, and the second- and third-week schedules identical to the previous cases. As a result, this protocol led to the lowest total tumor area among all the other simulated protocols, as shown in Figure 5B. The final tumor size reached only 8% of the untreated tumor, and 25% of the experimental tumor from protocol VA1 (Table 3).

### 3.4 Comparison of Optimized and Experimental Protocols

Of all the protocols that we investigated, the VA4 schedule was the most effective. The five daily vaccine injections in the first week (green plot, panel V of Figure 6) allowed for the early recruitment of T cells (green plot, panel T of Figure 6) by transfecting tumor cells in large amounts (compare green plots in panel I vs. U in Figure 6). Moreover, the addition of two doses of the anti-PD1 treatment earlier in VA4 than in the experimental protocol VA1, ensured better T cell activity and less exhaustion (compare green and black plots in panels T and J in Figure 6). The presence and activity of T cells in the first week were able to control the tumor size increase during this initial time period. This is in striking difference to the experimental protocol, in which the amount of untransfected cells steadily increased in size (black plot, panel U of Figure 6). In both cases, the amount of dead cells was similar in this early stage of treatment (panel W of Figure 6), showing that a reduction in tumor cell proliferation rather than tumor cell death was responsible for this observed effect. Over the first weekend (treatment holiday), T cells became inhibited; thus, there was a decline in the T plot and a rise in the J plot in Figure 6 (green lines). However, the second week started with two anti-PD1 injections (on Monday and Wednesday) to reactivate the inhibited T cells and to protect the free ones. This resulted in an increase in active T cells and a decrease in exhausted T cells over that time (green lines, panels T and J of Figure 6). The last week in protocol VA4 was very distinct from VA1. It consisted of two consecutive anti-PD1 injections at the beginning of the week (Monday and Tuesday; plot A in Figure 6) to reactivate the maximal number of T cells that were blocked during the weekend. This led to a significantly smaller total tumor area compared with the final tumor size after administration of the experimental protocol. Note that in the final period of VA4 treatment, the amount of dead cells was lower than in the VA1 protocol, but this was because the overall amount of untransfected and transfected tumor cells in VA4 was much lower than in VA1 (compare the green and black plots in panels W, U, and I of Figure 6). Among all the simulated protocols, the VA4 schedule resulted in the lowest total tumor area (Figure 5B).

**Figure 6:**
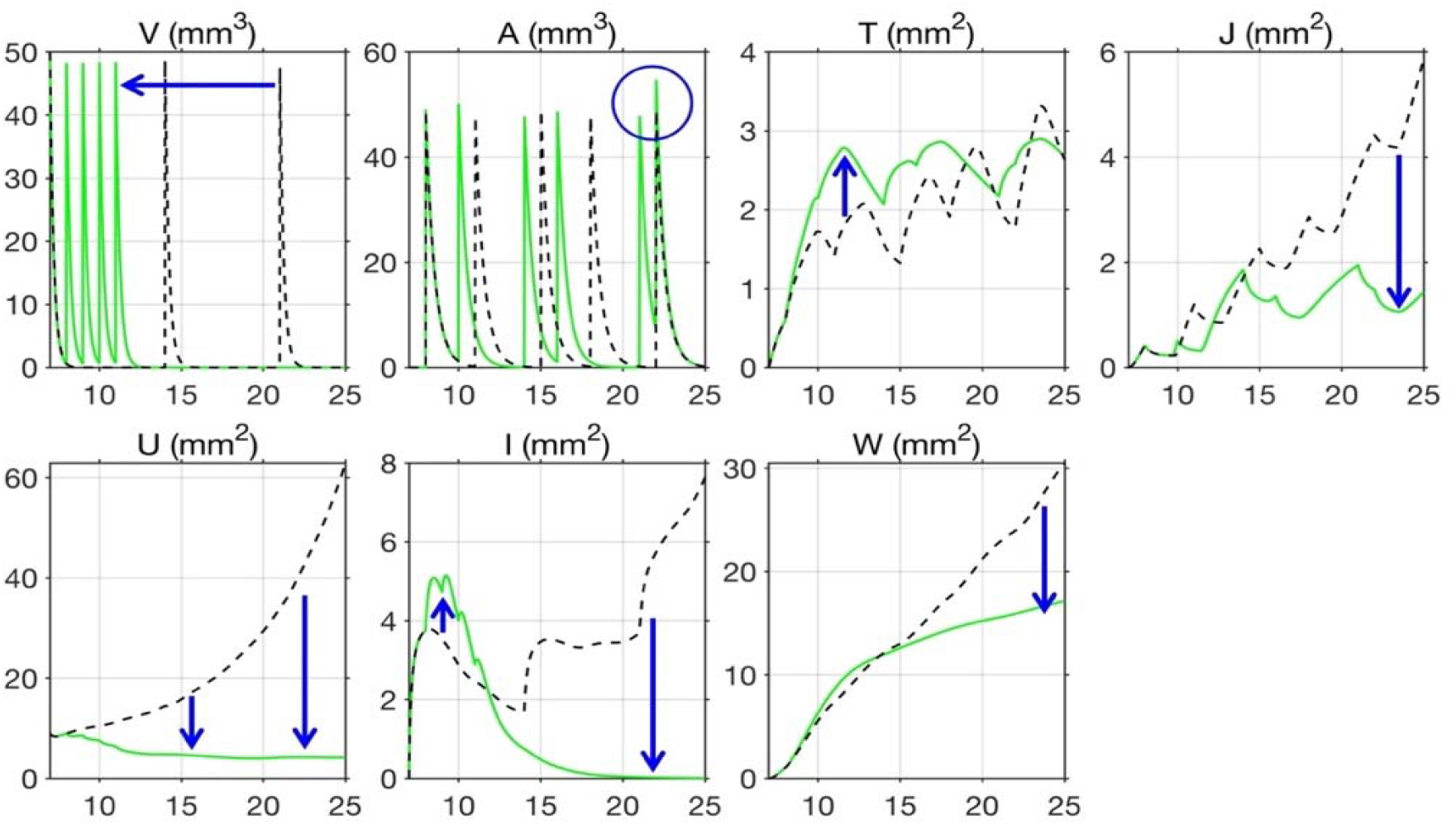
Comparison of optimized and experimental combination protocols. Each panel shows the time evolution of a different population of cells or treatments: V: vaccine, A: anti-PD1 treatment, T: active T cells, J: exhausted T cells, U: untrasfected tumor cells, I: transfected tumor cells, W: dead tumor cells. The course of an optimized protocol VA4 is shown in green, and experimental protocol VA1 in black. The blue circle in panel A highlights two consecutive injections in protocol VA4. The blue arrows represent important differences induced by the VA4 protocol leading to a reduced total tumor area.

In summary, our simulations suggest that injecting multiple doses of vaccine in the first week instead of spreading the injections over the entire three weeks will be beneficial in reducing tumor size. Our study also identified that keeping the same weekly schedule for anti-PD1 injections is not optimal. Instead, our recommendations for anti-PD1 injections are to administered them a day after vaccine injection in the first week to allow T cells to adequately infiltrate the tumor, and early in weeks 2 and 3 to maximize T cell reactivation.

## 4. Discussion

In this paper, we developed an ordinary differential equation model of a combination therapy consisting of a therapeutic anti-cancer emm55 vaccine and an anti-PD1 inhibitor. This model was calibrated to experimental data from a preclinical model of melanoma showing a time evolution of the tumor growth with and without the treatment. This model calibration process was performed in a three-stage hierarchical way by progressively fitting data from untreated tumors, tumors treated with the vaccine, and tumors treated with both the vaccine and checkpoint inhibitor. This allowed us to reduce the number of parameters that were fitted in each case by keeping fixed those that had already been calibrated in the previous model. Parameter fitting was performed using the fast MADS optimization method with rigorous convergence properties. Once calibrated, the model was used to determine the optimal treatment protocols for vaccine monotherapy and for the combined therapy of the vaccine and checkpoint inhibitor. As such, we were able to generate experimentally testable hypotheses for treatment scheduling. In the future, this method can be adjusted to model other vaccines and combination therapies, and can be applied to other cancers.

The goal of therapeutic cancer vaccines is to stimulate the patient’s immune system by providing priming antigens for the induction of a tumor-specific T cell response [3, 4]. The aim is to increase T cell infiltration into the tumor and to enhance T cell activity that may result in the eradication of minimal residual disease or in control of the growth of larger tumors, such as those discussed in this paper. However, the elevated numbers of tumor-reactive lymphocytes within the tumor tissue are also a desired feature in the novel immuno-oncology treatment—adoptive cell therapy (ACT) [43-45]. In this approach, the first step is to resect the patient’s tumor and expand tumor-infiltrating lymphocytes (TIL). Next, TIL that are reactive to the tumor are rapidly expanded and, finally, reinfused into the patient. If T cell infiltration into the tumor before its resection can be enhanced, the next steps of ACT will have a better chance of success. Therapeutic cancer vaccines may play this role. The emm55 vaccine discussed in this paper has shown anti-tumor efficacy toward melanoma in preclinical murine [18] and equine [46] studies. Moreover, the IL injection of the emm55-based vaccine was shown to be a feasible procedure in patients with metastatic melanoma in a recent clinical trial [47], which warrants further clinical studies. In these studies, well-designed treatment protocols with optimized drug administration schedules will be required to maximize treatment efficacy and the computational methods, like those discussed in this paper, may provide fast personalized approach to predict such protocols.

## Acknowledgments

This research was funded in part by the Moffitt Center for Immunization and Infection Research in Cancer (CIIRC) Award. The work was supported in part by the R01-CA259387 grant from the US National Institutes of Health, National Cancer Institute. The work of PB and MK was supported by the Jacobson Foundation for the Moffitt High-School Internship Program in Integrated Mathematical Oncology (HIP-IMO). This work was supported by the Shared Resources at the H. Lee Moffitt Cancer Center & Research Institute an NCI designated Comprehensive Cancer Center under the grant P30-CA076292 from the National Institutes of Health.

